# SimpleFold-Turbo: Adaptive Inference Caching Yields 14-fold Acceleration of Flow-matching Protein Structure Prediction

**DOI:** 10.64898/2026.04.07.714835

**Authors:** Geoffrey Taghon

## Abstract

We apply TeaCache, an adaptive caching technique from video diffusion to SimpleFold’s flow-matching protein structure prediction and achieve (9 to 14)-fold inference speedups with negligible quality loss. We determine that flow matching’s near-linear generative trajectories make consecutive neural-network evaluations highly redundant. At a low redundancy threshold, SimpleFold-Turbo (SF-T) skips ≈ 93 % of forward passes while preserving near-baseline template modeling (TM)-scores across 300 structurally diverse CATH domains and all six SimpleFold model sizes (100 million to 3 billion parameters), at compute budgets where log-uniform step-skipping collapses. Speedup scales with model size because caching overhead is constant while per-step cost grows, and a general three-phase skip pattern emerges independent of protein size or fold. SF-T requires no retraining, no weight modification, and no MSA server dependencies. We release SF-T as fully open-source software enabling thousands of structure predictions per hour on commodity hardware.

## 1 Introduction

### 1.1 The democratization gap

While protein structure prediction has been revolutionized by deep learning approaches, computational barriers remain substantial. A robust AlphaFold 3 inference pipeline requires high-end GPUs, typically available only as professional accelerators costing tens of thousands of dollars [1]. Structures for proteins exceeding 3,000 residues demand > 60 GB of GPU memory, and multiple sequence alignment (MSA)-dependent pipelines require multi-terabyte alignment databases regardless of query protein length. Some single-sequence models like ESMFold can predict structures without alignments [2], but GPU memory and compute time remain bottlenecks for high-throughput applications. Drug discovery campaigns routinely generate millions of sequence variants that could be binned and prioritized by predicted structure. Currently, resource-limited labs often lack this level of access on shared clusters or paid cloud instances [3, 4]. Judging by the state of open-source optimization techniques for image and 3D art models like StableDiffusion, “structure prediction for everyone” is likely possible but remains unfulfilled [5]. The caching opportunity Iterative generative models are often temporally redundant during successive compute steps. Multiple acceleration techniques exploit this redundancy by caching and reusing intermediate computations and skipping redundant steps. For example, TeaCache (timestep embedding-aware cache) uses inter-step embedding distance as a computationally cheap proxy for output similarity to greatly accelerate video generation [6]. At each generation step (video frame), TeaCache compares the current embedding to the previous; when the distance falls below a threshold, the expensive forward pass is skipped and the output is linearly interpolated from the last computed step. Video TeaCache achieves (2 to 5)-fold inference speedups with negligible quality loss. Similar strategies have been widely adopted in real-time rendering, such as NVIDIA’s DLSS, which runs at a reduced base rate and interpolates intermediate frames using motion vectors [7].

### 1.2 Why flow-matching is an ideal caching target

Caching is most effective when model outputs change smoothly between consecutive timesteps. Flow matching models predict a velocity field that transforms a simple distribution (typically Gaussian noise) into the data distribution (atomic coordinates) along an approximately linear trajectory [8]. Theoretically, if a straight line trajectory is learned, inference can be performed in a single step. In practice, multiple steps are used to correct for model imperfections, but the near linearity means that consecutive steps produce highly correlated outputs. This is precisely the redundancy that TeaCache exploits. Apple’s SimpleFold is a nearstate-of-the-art flow-matching model for single chain protein structure prediction that operates on atomic coordinates using SE(3)-equivariant transformer blocks [9]. The SimpleFold architecture requires no MSA computation and has fully open weights, making it an ideal test case for exploring caching-based acceleration. In the original benchmarks, protein modeling accuracy is comparable to AlphaFold2; mean TM-score performance versus ground truth reaches 72 to 99 % of AlphaFold2 baseline depending on dataset and SimpleFold model size. Importantly, standard SimpleFold on a standalone machine (64 GB MacBook Pro) reaches and, for sequences over 1024 amino acids, surpasses the inference speed of AlphaFold2 in a cluster setting (80 GB NVIDIA H100) [9].

### 1.3 Our contribution

Here we adapt TeaCache for SimpleFold to create SimpleFold-Turbo (SF-T). At each generative step, SF-T compares a scale-normalized input signal to the previous step; when the accumulated change remains below a threshold *τ*, the forward pass is skipped and the previous output is reused. This is an instant modification: no retraining, no weight changes, and no schedule tuning are required. We benchmark SF-T across all six SimpleFold model sizes on 300 structurally diverse CATH domains and release the full source code, which generates thousands of structures per hour on commodity hardware without discrete GPU, API, or MSA server dependencies (see § 4).

## 2 Results

### 2.1 Caching yields (9 to 14)-fold speedup with sub-angstrom fidelity

We evaluated SF-T across six SimpleFold model sizes (100M, 360M, 700M, 1.1B, 1.6B, and 3B parameters) on 300 structurally diverse CATH domains. At *τ* = 0.1, all models achieved approximately 93 % cache hit rates, computing a mean of 36 forward passes out of 500 (IQR: 34 to 37). Speedups scaled with model size: ≈ 9-fold for the 100M model, increasing to ≈ 14-fold for the 3B model (Figure 1A). This scaling occurs because caching overhead (computing embedding distances, interpolating outputs) remains constant while forward-pass cost grows with parameter count. The mean RMSD between SF-T and uncached SimpleFold predictions was 0.36 Å, below the ≈ 1.5 Å resolution typical of X-ray structures, indicating that caching introduces negligible coordinate perturbation relative to the model’s own accuracy (Figure 1A-B). Speedup also correlated positively with sequence length for all model sizes (*r* = 0.71 to 0.78, *p* < 0.001): Proteins of 300 to 450 residues achieved ≈ 11-fold mean speedup versus ≈ 8-fold for proteins under 100 residues, consistent with longer sequences having more cacheable trajectories.

**Figure 1.**
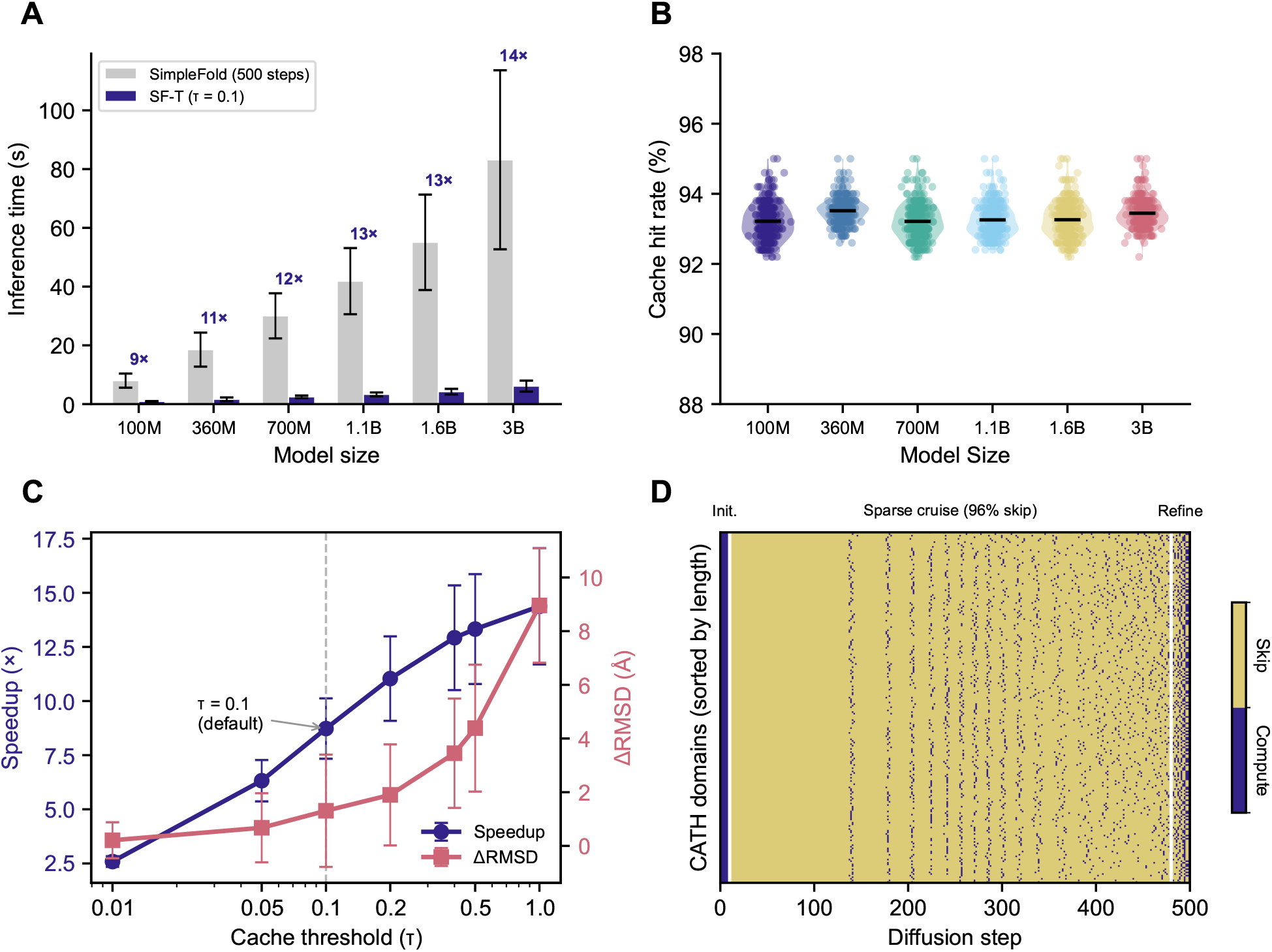
SF-T Performance and Mechanism. **(A)** End-to-end mean inference time per protein for SimpleFold (gray, 500 steps) versus SF-T (blue, *τ* = 0.1) across six model sizes, with fold-speedup annotated. Timings include ESM2-3B feature extraction and coordinate generation on Apple Mac Studio (M2 Max, 64 GB). **(B)** Mean cache hit rate remains constant at ≈ 93 % across model sizes. **(C)** Threshold (*τ*) sweep showing speed-quality tradeoff for SF-T-100M; *τ* = 0.1 achieves 9-fold speedup at 0.36 Å ΔRMSD. **(D)** Skip pattern heatmap (300 CATH domains × 500 steps) for SF-T-100M, revealing three-phase structure: initialization (steps 1 to 10, 0 % skip), sparse cruise (steps 11 to 480, 96 % skip), and refinement (steps 481 to 500, 64 % skip). The cruise-to-refinement transition may partly reflect SimpleFold’s logarithmic noise schedule. Violin plot means are indicated with horizontal bars. Error bars indicate

### 2.2 Threshold *τ* = 0.1 optimally balances speed and fidelity

To characterize the speed-to-quality tradeoff, we swept *τ* from 0.01 (highly conservative, nearly every step computed) to 1.0 (maximally aggressive, most steps skipped). At the conservative end (*τ* = 0.01), SF-T achieved only 2.6-fold speedup with 64 % cache hits, but maintained near-perfect coordinate fidelity (0.06 Å ΔRMSD vs. uncached). Increasing to *τ* = 0.05 yielded 6.5-fold speedup (88 % hits, 0.21 Å ΔRMSD). Our chosen default *τ* = 0.1 reached ≈ 9-fold speedup (93 % hits, 0.36 Å ΔRMSD), while more aggressive (*τ >* 0.2) traded fidelity for speed. We selected *τ* = 0.1 as the default because it maintains ΔRMSD below 0.5 Å while achieving roughly an order of magnitude acceleration (Figure 1C). Note that ΔRMSD here measures the relative structural deviation between cached and uncached predictions, not deviation from experimental ground truth.

### 2.3 A universal three-phase skip pattern emerges

To examine the mechanism of acceleration, we investigated the per-step decisions enforced by adaptive caching across the CATH subset. The following analyses used the smallest SimpleFold/SF-T variant (100M parameters, 386 MB) to provide a conservative quality baseline. When paired with the ESM2-3B encoder (quantized to 8-bits), this configuration represents the practical floor for edge deployment (≈ 4 GB total memory usage) [2]. Plotting the binary skip/compute decision for each of 500 steps across all 300 proteins revealed a striking pattern (Figure 1D). During steps 1 to 10, no steps are ever skipped; the network must establish initial coordinates from the input sequence in this phase. From step 11 onward, skip rates jump to 96 % through step 480. This is consistent with the near-linear trajectory assumption of flow matching: once the model has established the general direction of the generative path, the velocity field changes slowly enough that cached reuse suffices for the vast majority of steps. In the final 20 steps (481 to 500), skip rates drop to 64 %, reflecting elevated cache miss rates as the structure settles into its final conformation. The three-phase pattern (initialization, sparse cruise, refinement) was consistent across all proteins in our benchmark regardless of size or fold type.

### 2.4 Chain length predicts cacheability; fold type does not

Next, we asked what protein properties might determine cache efficiency. *K* -means clustering of proteins by their 500-step skip patterns yielded three groups with statistically identical secondary structure distributions (Fig. S1). Skip rate correlated strongly with sequence length (*r* = 0.78, *p* < 0.001): longer proteins (up to 446 residues in our dataset) achieved higher cache hit rates than shorter ones (minimum 50 residues). In contrast, secondary structure composition showed weak correlation with skip rate (*r* < 0.3). We also observed negligible correlation between skip rate and other protein properties like disorder, entropy, and hydrophobicity (Fig. S2). We attribute this length-cacheability relationship to the curse of dimensionality (see § 3). The absence of correlation with secondary structure or other biophysical properties supports this geometric interpretation (Figure 2A-B).

**Figure 2.**
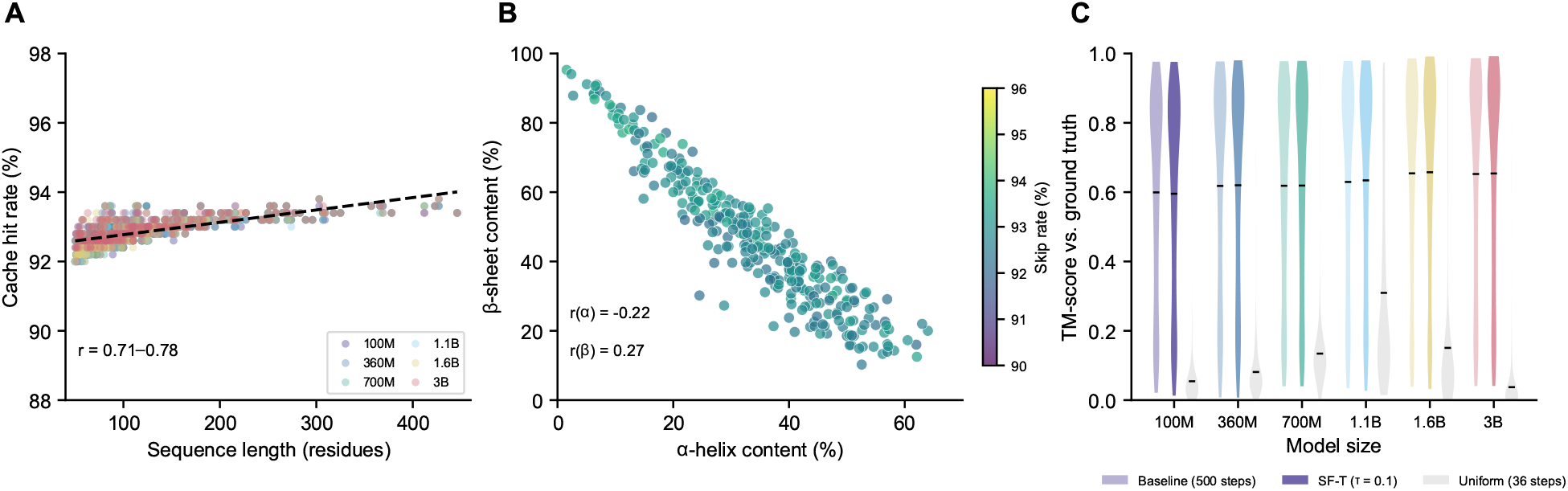
Cacheability Correlates and Quality Preservation. **(A)** Skip rate versus sequence length for all six models, (*r* = 0.71 to 0.78, *p* < 0.001), demonstrating that longer chains are more cacheable. **(B)** Skip rate versus secondary structure content shows negligible correlation (|*r* | < 0.3), indicating that fold composition does not determine cacheability. **(C)** Quality preservation: TM-score distributions for baseline SimpleFold, SF-T (*τ* = 0.1), and log-uniform step-skipping (36 steps, matching SF-T’s ≈36 computed steps at *τ* = 0.1) across all six model sizes, evaluated against PDB experimental structures. Violin plot means are indicated with horizontal bars, *n* = 300.

### 2.5 Quality is preserved against experimental structures

To ensure caching does not degrade biologically meaningful accuracy, we compared predictions from all six model sizes against CATH ground truth structures using template modeling (TM)-score and alpha carbon RMSD. We verified that our CATH benchmark domains were not present in SimpleFold’s training set by checking sequence identity against the published training splits. Across all models, baseline SimpleFold (500 steps, no caching) achieved mean TM-scores ranging from 0.599 (100M; 95 % bootstrap CI: 0.567 to 0.632) to 0.654 (1.6B). SF-T at *τ* = 0.1 matched baseline quality at every model size: TM-scores ranged from 0.595 (100M; CI: 0.562 to 0.628) to 0.657 (1.6B). The mean ΔTM-score was -0.004 (CI: -0.051 to +0.042) for the 100M model and +0.002 (CI: -0.046 to +0.050) for the 3B model, with confidence intervals spanning zero in both cases (Figure 2C, Fig. S3).

### 2.6 Adaptive caching vastly outperforms static step reduction

To provide a standard baseline for diffusion model acceleration, we swept log-uniform step-skipping across nine step counts (12 to 250) for all six model sizes, matching SimpleFold’s native logarithmic noise schedule (Fig. S4). At matched compute budgets (approximately 36 computed steps), adaptive caching achieved TM-scores of 0.595 to 0.658 across models, while log-uniform step-skipping with 36 log-spaced steps collapsed to TM-scores of 0.037 to 0.309, a difference of +0.33 to +0.62 in favor of adaptive caching. Log-uniform scheduling required approximately 100 steps (mean TM-score = 0.611) to approach the quality that adaptive caching achieved with 36. Below 59 log-uniform steps, predictions degraded severely (mean TM-score = 0.438); below 24 steps, they were structurally meaningless (mean RMSD > 97 Å). This pattern was consistent across all model sizes and sequences, confirming that adaptive caching’s advantage is not protein- or model-specific but reflects a fundamental property of how compute should be allocated along a flow-matching trajectory (Fig. S4).

## 3 Discussion

### 3.1 Adaptive caching as adaptive integration

Adaptive caching can be understood as a form of adaptive step-size selection. In a flow-matching model, the generation process integrates a learned velocity field over *T* discrete steps. When the velocity field is locally smooth, large integration steps introduce minimal error and multiple small steps are redundant. TeaCache detects these smooth regions via embedding distance and effectively increases the integration step size by reusing the previous output rather than recomputing. Viewed this way, SF-T’s ≈ 93 % cache hit rate implies that SimpleFold’s 500-step schedule is approximately 10-fold oversampled for most of the trajectory. Critically, this oversampling cannot be resolved by simply reducing the step count: log-uniform scheduling requires ≈ 100 steps to approach the quality that adaptive caching achieves with 36 (Fig. S4), confirming that SF-T concentrates compute on the steps where the trajectory actually changes direction.

### 3.2 Flow matching explains extreme cacheability

Why does SF-T achieve 14-fold speedup via caching while typical video diffusion caching manages only ≈ 4-fold? We attribute this primarily to flow matching’s near-linear trajectories, which produce highly correlated consecutive outputs. Cached reuse of previous step outputs is a reliable approximation because the velocity field changes minimally between adjacent steps. Static step-skipping, by contrast, removes intermediate steps entirely rather than evaluating whether their outputs have materially changed. The three-phase skip pattern we observe is consistent with this explanation: the initialization phase establishes the trajectory direction, the cruise phase traverses the near-linear middle segment, and the refinement phase shows elevated cache miss rates, suggesting the velocity field changes more rapidly as the prediction converges.

The strong correlation between chain length and cache efficiency (*r* = 0.78) admits a straightforward mathematical explanation: the curse of dimensionality. As chain length increases, the coordinate representation grows in dimensionality, and consecutive embeddings tend to be closer in relative L1 distance, causing TeaCache to mark more steps as skippable. To test whether dimensionality alone accounts for this relationship, we constructed a control experiment with synthetic flat random vectors matching the dimensionality distribution of our protein set. The synthetic vectors showed a weaker length-cacheability correlation (*r* = 0.46) compared to actual protein trajectories (*r* = 0.78), indicating that while dimensionality contributes, the structured nature of protein generative trajectories amplifies the effect beyond what geometry alone predicts (Fig. S6). Secondary structure composition, disorder propensity, hydrophobicity, and atom distance from backbone all showed negligible correlation with cacheability (|*r* | < 0.3; Figs. S2, S5), further confirming that skip patterns reflect trajectory geometry rather than biophysical properties.

### 3.3 Practical implications

SF-T’s order-of-magnitude acceleration has immediate practical consequences. A 3B parameter model running at 14-fold speedup outperforms a 360M model at baseline speed, enabling deployment of large models on accessible hardware (e.g., Apple Silicon, consumer GPU lines). Furthermore, like SimpleFold, SF-T has no MSA or template server dependencies and thus can operate on machines disconnected from the Internet. High-throughput applications such as screening millions of sequence variants for drug discovery become feasible on modest, air-gapped GPU clusters. Sub-second coordinate predictions enable interactive structural hypothesis testing during experimental design. Where compute energy and carbon footprint matter, SF-T delivers equivalent accuracy with a 93 % reduction in compute.

### 3.4 Generalizability and limitations

SimpleFold was chosen as the initial test case because it is, to our knowledge, the only near-state-of-the-art protein structure prediction model built on a flow-matching architecture with fully open weights, a public codebase, and no MSA or internet dependencies. Because our results suggest that extreme cacheability is a property of flow-matching trajectories rather than a domain-specific phenomenon, we predict that any flow-matching generative model should exhibit similar speed gains from TeaCache-style adaptive caching. As flow matching is increasingly adopted for biomolecular modeling, including RNA structure, small molecule conformers, and protein complexes, this technique should transfer directly with no retraining required. We use the MLX backend on Apple Silicon for maximum inference speed but also provide fallback to PyTorch for compatibility with CPUs and CUDA. We quantized ESM2-3B to 8 bits after determining that feature outputs were unaffected compared to full 32-bit parameters.

This work has several limitations. We evaluated only the SimpleFold architecture family with single protein chains; testing on other flow-matching models (e.g., Boltz [10]) is an important next step. While detailed skip-pattern analysis used the 100M model, the log-uniform-versus-adaptive comparison was performed across all model sizes (100M to 3B), confirming that the adaptive advantage is model-size independent (Fig. S4). We note that more sophisticated baselines are possible: an adaptive ODE solver rather than the current Euler-Maruyama solver or a hand-crafted endpoint-heavy schedule (concentrating steps where the trajectory curves most) could narrow the gap between log-uniform and adaptive caching, though the simplicity and generality of the training-free TeaCache approach remains attractive. For MSA-reliant architectures such as AlphaFold 3, the MSA computation itself remains a bottleneck (≈ 10 s) that caching does not address [11]. The optimal cache *τ* may vary by application: users prioritizing throughput might prefer *τ* = 0.2, while those requiring maximum accuracy could use *τ* = 0.05.

## 4 Methods

### 4.1 Dataset

We benchmarked SF-T versus SimpleFold on 300 structurally diverse domains from the CATH database [12], selected to span all major fold classes with sequence lengths ranging from 50 to 446 residues (“CATH-300”). We used a greedy selection algorithm to maximize structural diversity. One domain (1hkgA03) was replaced due to missing residues in the ground truth PDB; 1byuB00 was substituted at identical sequence length. All 300 domains had valid ground truth structures for quality evaluation. Generated structure files for CATH-300 using all SimpleFold/SF-T model sizes are available upon request.

### 4.2 Models

SimpleFold is a flow-matching protein structure prediction model using general-purpose transformer blocks with adaptive layers [9]. We evaluated all six model sizes (100M, 360M, 700M, 1.1B, 1.6B, 3B parameters), each running 500 diffusion steps with a logarithmic noise schedule. The architecture is SE(3)-equivariant, operating directly on atomic coordinates.

### 4.3 SF-T caching procedure

SF-T adapts the timestep embedding-aware cache (TeaCache) algorithm [6] for protein structure generation. At each step *t ∈* [0, 1], we compute a scalar modulated input

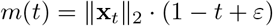

where **x**_*t*_ are the current noisy coordinates and *ε* = 0.1 prevents numerical instability near *t* = 1. We maintain a running accumulated relative difference:

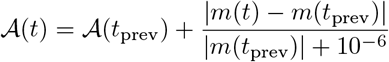

At each step, the cache decision is first evaluated, the accumulator reset if needed, and the velocity assigned:

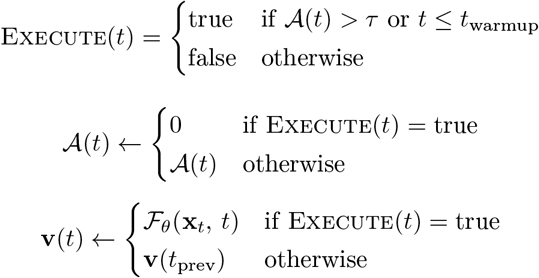

The first *t*_warmup_ = 10 steps are always computed to establish the trajectory direction. The cache decision is made per-sample (batch size = 1 throughout).The default *τ* = 0.1 was selected to maintain ΔRMSD *<* 0.5 Å on the CATH-300 structure dataset.

### 4.4 Analysis

Skip patterns were analyzed by extracting the binary skip/compute decision at each of 500 steps for each protein. *K* -means clustering (*k* = 3) of skip patterns was performed directly on binary skip masks, with cluster count selected to balance interpretability and statistical separation. Structure quality was assessed using TM-score, RMSD, and lDDT computed against corresponding experimental ground truth structures from the CATH dataset. Correlation between skip rate and protein properties used Pearson’s *r* with a significance threshold *p* < 0.001. Secondary structure fractions were computed from *ϕ*/*ψ* backbone dihedral angles using standard Ramachandran regions: helix (-100° < *ϕ* < -30°, -80° < *ψ* < -5°), sheet (-180° < *ϕ* < -30°, 50° < *ψ* < 180°), with remaining residues assigned to coil. Sequence properties were computed as follows: hydrophobicity used the Kyte-Doolittle scale averaged over all residues; disorder propensity was the fraction of disorder-prone residues (P, S, G, Q, N, K, D, E, R); sequence entropy was Shannon entropy over the amino acid frequency distribution.

### 4.5 Hardware

All computation was performed on an Apple Mac Studio (Model Z17Z000JXLL/A, 2023) with an M2 Max processor and 64 GB of memory.

### 4.6 Software and Data

SF-T source code, evaluation scripts, and the CATH-300 domain list, are available on [GitHub](https://github.com/usnistgov/simplefold-turbo). The codebase is built on SimpleFold [9] using the MLX backend, with PyTorch fallback. ESM2-3B was quantized to 8-bit precision using MLX’s built-in quantization and is available for download via Hugging Face Hub (gtaghon/esm3-sm-open-v1-mlx-8bit). All inference used deterministic seeding (seed = 42 + protein index). Claude (Anthropic) was used for code development and documentation. All raw data including 30 000+ structural models generated during this work is published at [Zenodo](https://zenodo.org/uploads/19455758) for inspection and analysis.

## 5 Acknowledgments

We thank Bruce Wittmann (Microsoft) and Zoila Jurado (NIST) for insightful feedback on the manuscript.

G.T. was supported in part by an appointment to a Research Associateship Program at the National Institute of Standards and Technology, administered by the Johns Hopkins University Whiting School of Engineering.

