## Supplementary Figures for "SimpleFold-Turbo: Adaptive Inference Caching Yields 14-fold Acceleration of Flow-matching Protein Structure Prediction"

**Geoffrey Taghon**[1,2,†]

[1]National Institute of Standards and Technology, Gaithersburg, MD, USA
[2]Whiting School of Engineering, Johns Hopkins University, Baltimore, MD, USA

**Date:** April 7, 2026 | **Code:** github.com/usnistgov/simplefold-turbo

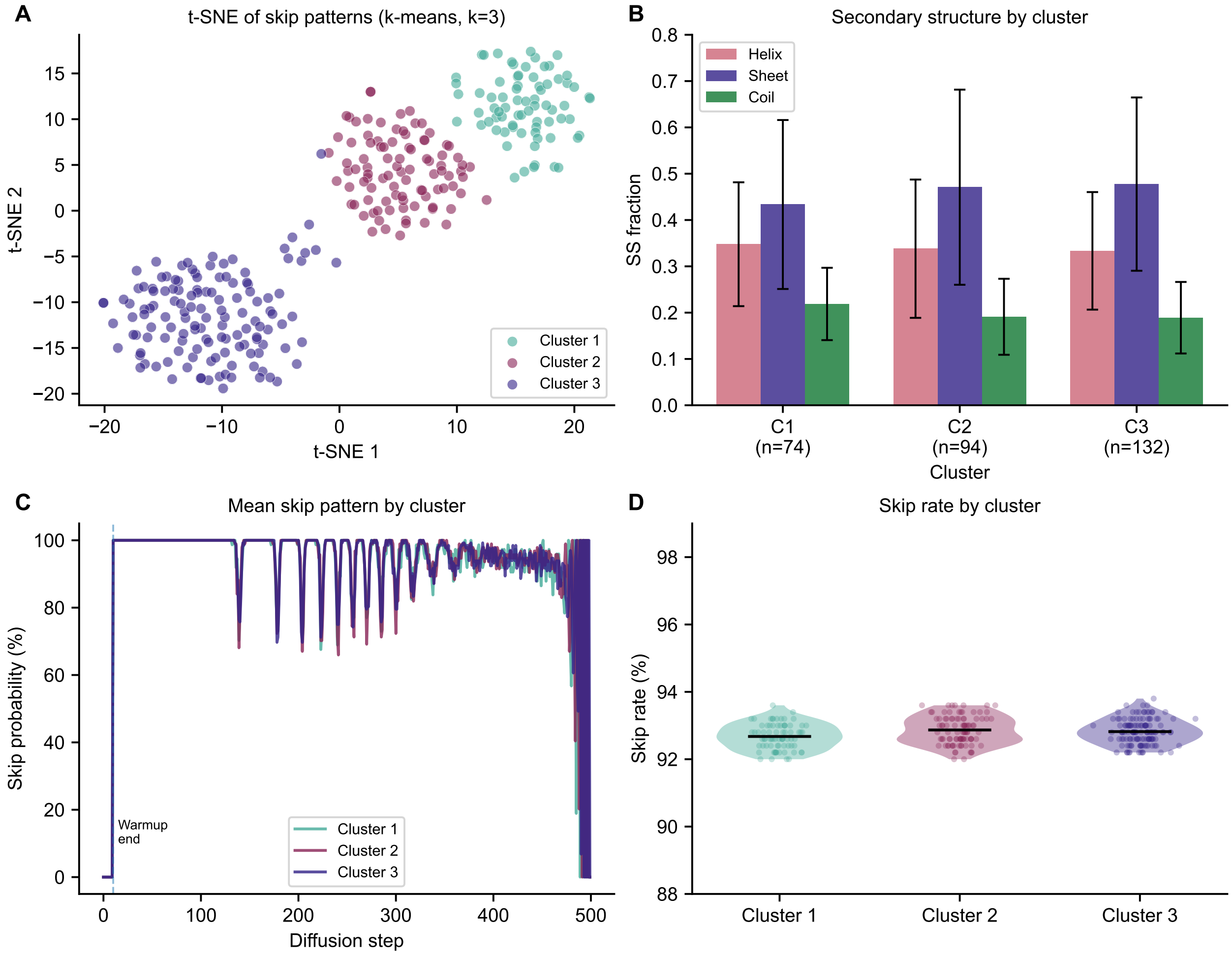


**Figure S1.** **Skip pattern is not dependent on amino acid properties (SF-T-100M).** Skip pattern is not dependent on amino acid properties (100M model). **(A)** $k$-means clustering ($k = 3$) reveals distinct skip pattern clusters, but **(B)** they contain an equivalent distribution of secondary structures. **(C)** All proteins tend to follow the same general skip pattern with visible minima that act as "keyframes" for interpolating between until the last third of the diffusion steps, where they diverge. **(D)** However, all clusters show statistically identical overall skip rate, indicating that the caching method should be generally applicable to diverse protein sequences. Violin plot means are indicated with horizontal bars. Error bars indicate $\pm$ 1 SD, $n = 300$.

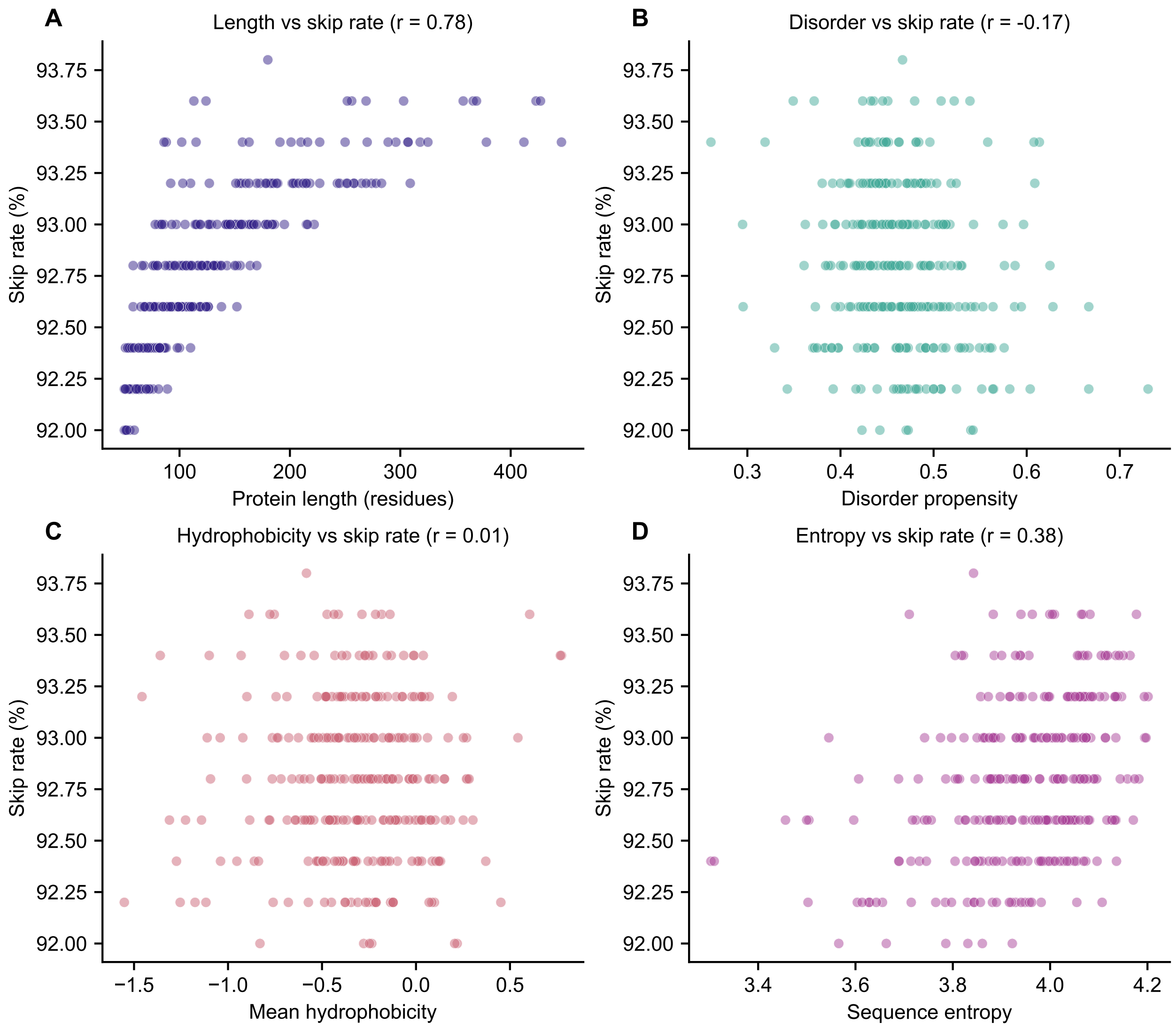


**Figure S2. Skip rate correlates with sequence length, but not other properties (SF-T-100M). (A)** Skip rate is strongly correlated ($r = 0.78$) to sequence length, but not other protein properties like **(B)** structural disorder or **(C)** hydrophobicity. **(D)** Entropy shows a moderate correlation with skip rate ($r = 0.38$), likely reflecting its covariance with sequence length.

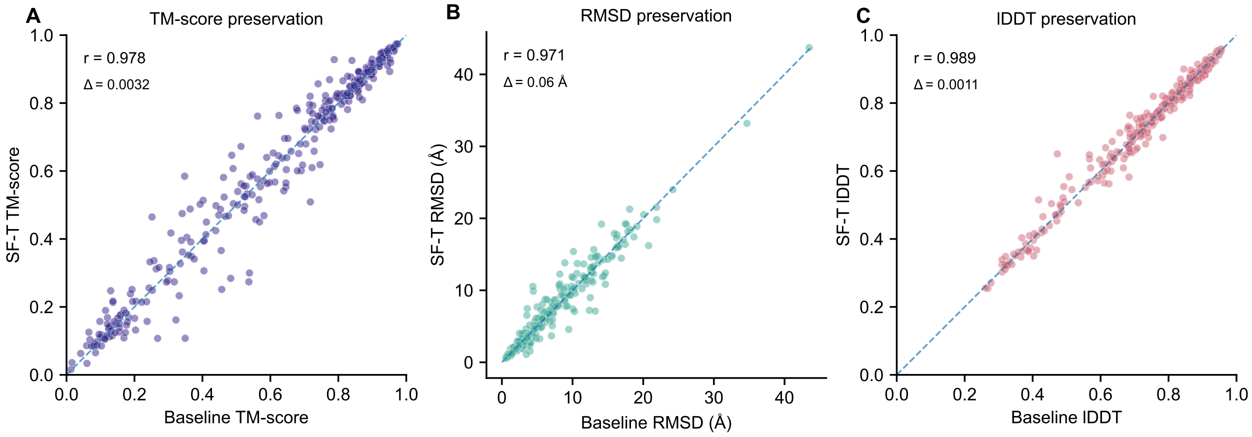


**Figure S3. Quality differences between uncached baseline (SimpleFold-100M) and cached (SF-T-100M) are negligible. (A)** TM-score, **(B)** RMSD, and **(C)** lDDT were highly correlated with (SF-T) and without (SimpleFold) adaptive caching ($n = 300$).

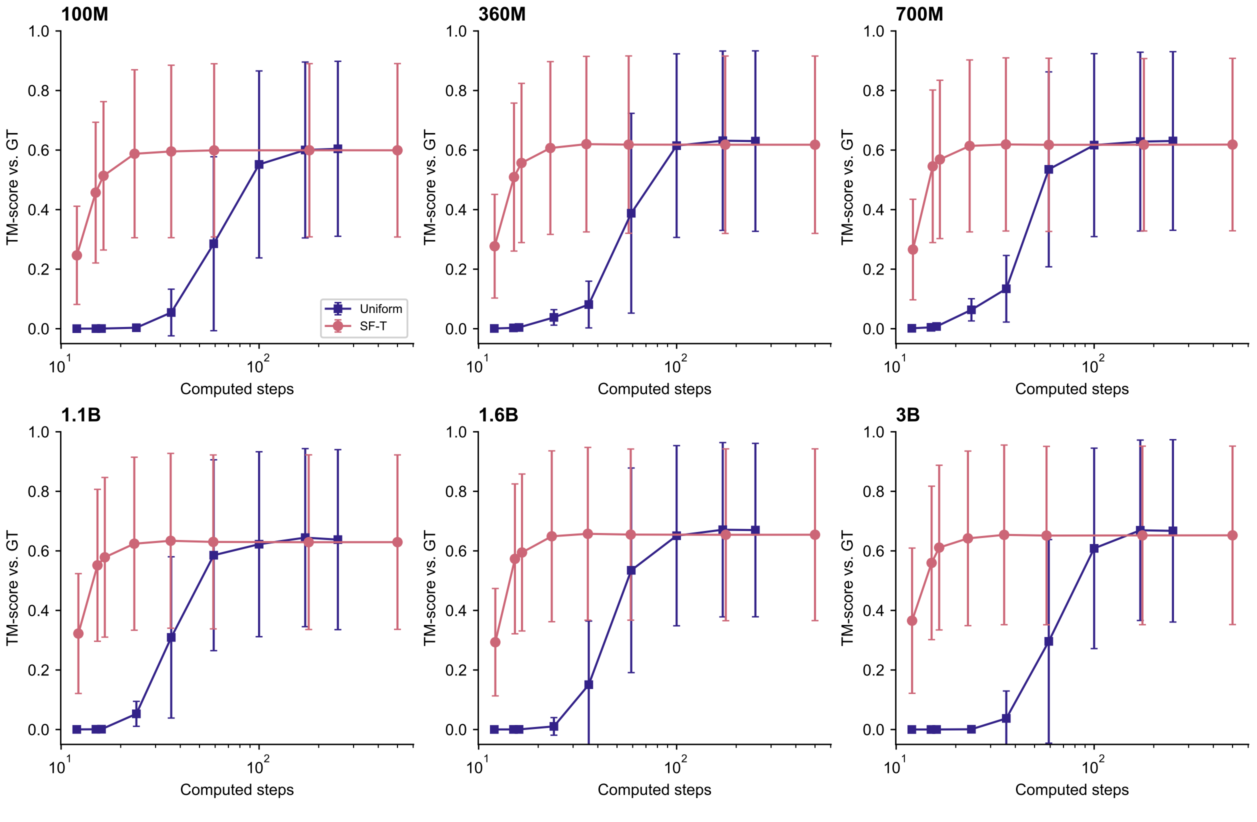


**Figure S4. Log-uniform step-skipping versus adaptive caching (SF-T) across all model sizes.** For each of six SimpleFold models, TM-score versus number of computed steps is shown for log-uniform step-skipping (9 step counts, 12 to 250) and adaptive caching (8 thresholds, $\tau = 0.0$ to 1.0). Adaptive caching preserves near-baseline quality at lower compute budgets while log-uniform step-skipping degrades catastrophically below $\approx 100$ steps. The pattern is consistent across all model sizes. Error bars indicate $\pm$ 1 SD, $n = 300$.

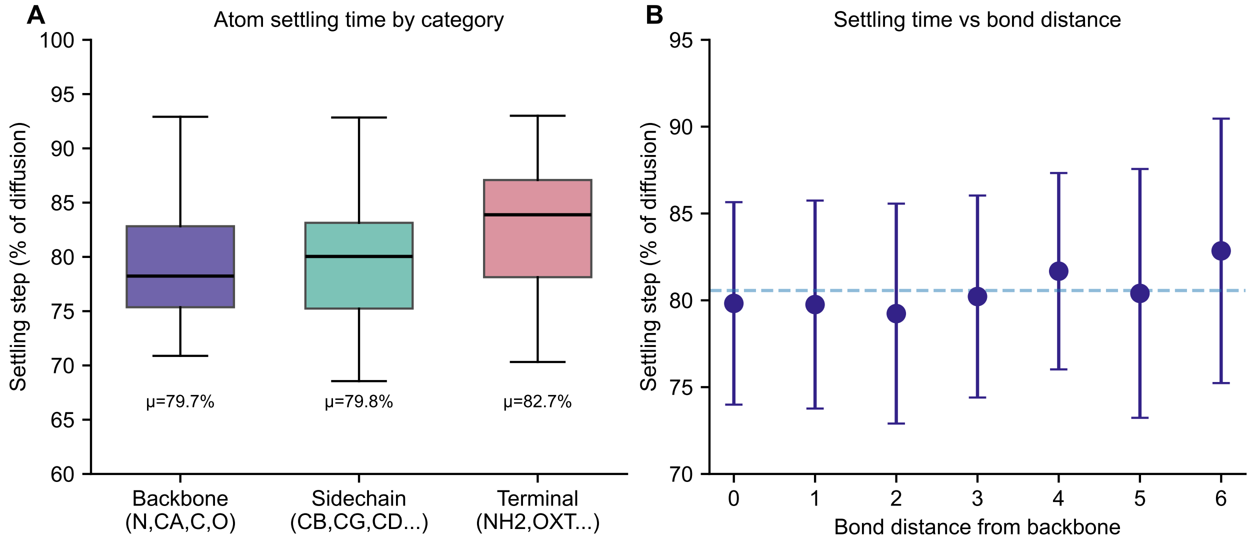


**Figure S5. Atom distance from backbone is not a factor during caching (SF-T-100M).** Contrary to our expectations, atoms further from the backbone did not have longer coordinate settling times than the backbone during the coordinate generation process. **(A)** Neither the relative atom group (backbone, sidechain, terminal) nor **(B)** explicit atom position in the amino acid had significant trends with respect to number of steps needed to settle into position. Error bars indicate $\pm$ 1 SD, $n = 300$.

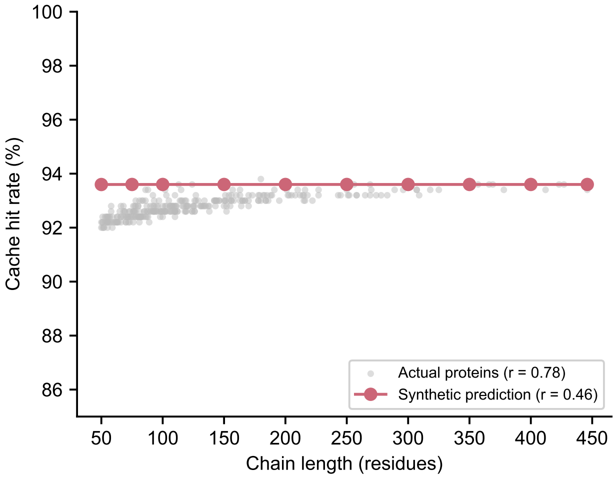


**Figure S6. Dimensionality control experiment (SF-T-100M)**. Comparison of length-to-cacheability correlation in actual protein trajectories ($r = 0.78$) versus synthetic flat random vectors matched in dimensionality ($r = 0.46$). The stronger correlation in real proteins indicates that structured generative trajectories amplify the dimensionality effect beyond pure geometric expectation.
